# Developmental Mechanisms Linking Form and Function During Jaw Evolution

**DOI:** 10.1101/264556

**Authors:** Katherine C. Woronowicz, Stephanie E. Gline, Safa T. Herfat, Aaron J. Fields, Richard A. Schneider

## Abstract

How does form arise during development and change during evolution? How does form relate to function, and what enables embryonic structures to presage their later use in adults? To address these questions, we leverage the distinct functional morphology of the jaw in duck, chick, and quail. In connection with their specialized mode of feeding, duck develop a secondary cartilage at the tendon insertion of their jaw adductor muscle on the mandible. An equivalent cartilage is absent in chick and quail. We hypothesize that species-specific jaw architecture and mechanical forces promote secondary cartilage in duck through the differential regulation of FGF and TGFβ signaling. First, we perform transplants between chick and duck embryos and demonstrate that the ability of neural crest mesenchyme (NCM) to direct the species-specific insertion of muscle and the formation of secondary cartilage depends upon the amount and spatial distribution of NCM-derived connective tissues. Second, we quantify motility and build finite element models of the jaw complex in duck and quail, which reveals a link between species-specific jaw architecture and the predicted mechanical force environment. Third, we investigate the extent to which mechanical load mediates FGF and TGFβ signaling in the duck jaw adductor insertion, and discover that both pathways are mechano-responsive and required for secondary cartilage formation. Additionally, we find that FGF and TGFβ signaling can also induce secondary cartilage in the absence of mechanical force or in the adductor insertion of quail embryos. Thus, our results provide novel insights on molecular, cellular, and biomechanical mechanisms that couple musculoskeletal form and function during development and evolution.

## Introduction

One of the most remarkable aspects of being an embryo, and a phenomenon that has intrigued embryologists since Aristotle, is the ability to grow in a manner “rather prospective than retrospective” (Thompson, 1942). In theory, how the form of an embryo can presage later adult function is explained by Aristotle’s observation that “the organism is the *τελoς*, or final cause, of its own process of generation and development” (Thompson, 1942). But elucidating precise molecular mechanisms that link form and function, and specifically whether form arises from function or function follows form remains challenging, because, like the chicken and the egg, form and function are seamlessly intertwined during development and evolution.

Some of the most illustrious instances of form and function appear in the craniofacial complex in birds, which are masters of adaptation. A specialized beak seems to exist for every avian diet: insectivore, granivore, nectarivore, frugivore, carnivore, omnivore, etc. (Schneider, 2007; Zusi, 1993). Each diet is supported by a range of structural adaptations to the jaw including size, shape, and sites of muscle attachments (Fish and Schneider, 2014b; Tokita and Schneider, 2009). For example, in *Anseriformes*, or waterfowl such as duck, which use their broad bills to dredge sediment for food, the mandibular adductor (MA) muscle attaches laterally to a large protruding coronoid process (CP) on the mandible. Such a configuration provides a robust insertion site for transmitting the high magnitude forces associated with suction pump and levered straining jaw movements (Dawson et al., 2011; Zweers, 1974; Zweers et al., 1977). In duck, as in humans, the CP develops via a secondary cartilage intermediate (Solem et al., 2011). Secondary cartilage requires proper mechanical stimulation for its induction and maintenance, as confirmed by explant cultures and paralysis experiments, and is a feature of many joints in neognathic avian skulls, as well as in select tendon and muscle insertions (Hall, 1967, 1968, 1972, 1986). In paralyzed duck, secondary cartilage fails to form at the CP, suggesting that the mechanical environment (i.e., function) during development promotes secondary chondrogenesis (Solem et al., 2011). By comparison, *Galliformes* like quail and chick, feed primarily by pecking seed, and this is reflected in the relatively gracile construction of the jaw and adductor muscles, which insert dorsally on the mandible and lack secondary cartilage on the CP. Exploiting such species-specific differences in quail and duck, as we have done previously in studies of beak, feather, cartilage, bone, and muscle patterning, (Ealba et al., 2015; Eames and Schneider, 2008; Fish and Schneider, 2014a; Hall et al., 2014; Schneider, 2005, 2015; Schneider and Helms, 2003; Tokita and Schneider, 2009) provides an opportunity to investigate molecular, cellular, and biomechanical mechanisms that integrate form and function in the jaw apparatus during development and evolution.

The species-specific jaw morphology that distinguishes duck from quail is mediated by the neural crest mesenchyme (NCM), which gives rise to all of the associated cartilage, bone, and muscle connective tissues (Noden and Schneider, 2006). Transplanting NCM from quail into duck has established that NCM controls the size and shape of the jaw skeleton, as well as the orientation and insertion of muscles (Ealba et al., 2015; Eames and Schneider, 2008; Fish and Schneider, 2014a; Hall et al., 2014; Schneider and Helms, 2003; Solem et al., 2011; Tokita and Schneider, 2009). Chimeric “quck” develop a quail-like jaw musculoskeleton including a dorsal MA insertion that lacks secondary cartilage. The precise developmental mechanisms through which this happens have remained an open question. Presumably, for such a transformation, quail NCM alters the duck-host environment in a manner that changes not only the form of the jaw apparatus but also the function, since the presence or absence of secondary cartilage depends upon proper mechanical cues. In this context, the lateral versus dorsal insertion of the MA muscle might produce distinct mechanical forces, but differences in the quantity and/or quality of such forces in quail versus duck are completely unknown. Furthermore, those signaling pathways that are mechanoresponsive and ultimately govern species-specific adaptation to the mechanical environment remain unclear. The current study set out to address these unresolved issues.

We hypothesized that the form of the duck MA complex creates a species-specific mechanical environment, which activates molecular programs for secondary chondrogenesis at the CP. To test our hypothesis, we employed a range of strategies. We modulated the form of the duck MA complex by titrating the amount of donor versus host NCM-derived tissues in chick-duck chimeras. We quantified embryonic jaw motility in duck versus quail and performed finite element analysis (FEA) to model the mechanical environment of the MA complex. We employed FEA in order to make predictions about the extent to which mechanical forces might underlie the induction of secondary cartilage and the differential regulation of mechanically responsive signaling pathways. We disrupted the mechanical environment of the MA complex by paralyzing duck embryos and then we assayed for changes in signaling pathways that might be mechanically responsive at the CP. After identifying candidate pathways, we tested if they were necessary and/or sufficient for the formation of secondary cartilage.

Our results reveal that the formation of secondary cartilage on the CP depends upon the amount and spatial distribution of NCM-derived connective tissues. While we observe few quantitative differences in the amount of motility between quail and duck, our FEA suggests that quail and duck have qualitatively distinct mechanical forces at the MA insertion. Additionally, we discover that both FGF and TGFβ signaling are responsive to mechanical forces within the duck MA complex, and are necessary for secondary chondrogenesis at the CP. Additionally, we find that exogenous FGF and TGFβ ligands can rescue cartilage in paralyzed duck and also induce cartilage in the quail MA insertion, where ordinarily there is none. Overall, this study provides mechanistic insights on how species-specific morphology, mechanical forces, and resultant changes in signaling activity become integrated and contribute to musculoskeletal plasticity. While form initially dictates function, function can also act as a potent modulator of musculoskeletal form during development and evolution.

## Methods

### The use of avian embryos

Fertilized eggs of Japanese quail (*Coturnix coturnix japonica*) and white Pekin duck (*Anas platyrhynchos*) were purchased from AA Lab Eggs (Westminster, CA) and incubated at 37.5°C in a humidified chamber (GQF Hova-Bator, Savannah, GA) until embryos reached stages appropriate for manipulations, treatments, and analyses. For all procedures, we adhered to accepted practices for the humane treatment of avian embryos as described in S3.4.4 of the AVMA Guidelines for the Euthanasia of Animals: 2013 Edition (Leary et al., 2013). Embryos were stage-matched using an approach that is based on external morphological characters and that is independent of body size and incubation time (Hamilton, 1965; Ricklefs and Starck, 1998; Starck and Ricklefs, 1998). The Hamburger and Hamilton (HH) staging system, originally devised for chick, is a well-established standard (Hamburger and Hamilton, 1951). Separate staging systems do exist for duck (Koecke, 1958) and quail (Ainsworth et al., 2010; Nakane and Tsudzuki, 1999; Padgett and Ivey, 1960; Zacchei, 1961) but these embryos can also be staged via the HH scheme used for chicken (Ainsworth et al., 2010; Le Douarin et al., 1996; Lwigale and Schneider, 2008; Mitgutsch et al., 2011; Schneider and Helms, 2003; Smith et al., 2015; Starck, 1989; Yamashita and Sohal, 1987; Young et al., 2014). Criteria utilized to align quail and duck at a particular HH stage change over time depending on which structures become prominent. For early embryonic stages, we used the extent of neurulation, NCM migration, and somitogenesis as markers (Fish et al., 2014; Lwigale and Schneider, 2008; Schneider and Helms, 2003); whereas later, we relied on growth of the limbs, facial primordia, feather buds, and eyes since these become more diagnostic (Eames and Schneider, 2005; Merrill et al., 2008).

### Histology

Embryos were fixed overnight in 10% neutral buffered formalin at 4°C, paraffin embedded, and sectioned at 10μm. Cartilage, bone, muscle, and tendon were visualized using Milligan’s Trichrome or Safranin-O (Ferguson et al., 1998; Presnell and Schreibman, 1997).

### Clearing and staining

Embryos were fixed overnight at 4°C in 10% neutral buffered formalin before clearing and staining with Alcian Blue and Alizarin Red to visualize cartilage and bone of the jaw complex including the CP (Wassersug, 1976).

### cDNA preparation

RNA was isolated from microdissected duck samples using the ARCTURUS PicoPure RNA Isolation Kit (ThermoFisher, Waltham, MA). Reaction specifications and reverse transcription programs were followed as previously published (Ealba and Schneider, 2013).

### In situ hybridization

Spatial and temporal patters of gene expression were analyzed by in situ hybridization as previously described (Albrecht et al., 1997; Schneider et al., 2001). Species-specific probes against duck FGF and TGFβ ligands (*Fgf4, Fgf8, Tgfβ2, Tgfβ3*), receptors (*Fgfr2, Fgfr3, Tgfβr2*), and downstream effectors (*Pea3, Erm, and Smad3*), were cloned from duck HH33 cDNA libraries isolated from whole heads (Table S1). Probes were designed to recognize all isoforms. High fidelity Pfu DNA polymerase (Strategene, La Jolla, CA) was used to amplify target genes. The protocol was: step 1, 2 minutes at 94°C; step 2, 30 seconds at 94°C; step 3, 30 seconds at 37.5°C; step 4, 2 minutes at 72°C; step 5, repeat steps 2 to 4 39 times; step 6, 5 minutes at 72°C; step 7, hold at 4°C. PCR products were run on a 1% agarose gel. Bands of the appropriate molecular weight were gel extracted using QIAEX II Gel Extraction Kit (Qiagen, Hilden, Germany). PCR products were ligated into pGEM-T Easy Vector System I (Promega, Madison, WI) or CloneJET PCR Cloning Kit (ThermoFisher, Waltham, MA) and used to transform NEB 5α E. coli cells (New England Biolabs, Ipswitch, MA). Clones were sequenced (McLab, South San Francisco, CA) using a T7 promoter primer. Sequencing results were analyzed using Geneious (Biomatters, Auckland, New Zealand). Once probe sequences were confirmed, DIG-labeled RNA probes were synthesized using DIG RNA labeling mix (Roche, Basel, Switzerland). Cloned species-specific duck probes were used to identify gene expression patterns in embedded and sectioned HH33 and HH36 paralyzed and stage matched control duck.

### TUNEL staining

10µm tissue sections of duck embryos 24 hours after treatment with SU5402, SB431542, or DMSO soaked beads were processed using a fluorescent TUNEL staining kit (Roche, Basel, Switzerland). As a positive control, DNase was added to a subset of DMSO-treated tissue sections. The percentage of cell death was quantified using 3D microscopy processing software Imaris (Bitplane, Belfast, United Kingdom). Image intensity was rendered in 3D and Hoescht (Sigma-Aldrich, St. Louis, MO) and TUNEL-stained nuclei within 100μm of the implanted bead were counted using software-enabled volumetric criteria (surface detail=5μm, background subtraction=12μm, seed point diameter=30μm). Statistical significance was determined by ordinary one-way ANOVA (Prism 7, GraphPad Software, Inc., La Jolla, CA).

### Surgical bead implantation

10mM of SU5402 (Sigma-Aldrich, St. Louis, MO), a small molecule that prevents autophosphorylation of receptor tyrosine kinases and is most specific to FGFRs (Sun et al., 1999; Sun et al., 1998), and 100mM of SB431542 (Santa Cruz Biotechnology, Santa Cruz, CA), a small molecule that inhibits autophosphorylation of TGFβRs (Callahan et al., 2002; Inman et al., 2002), were diluted in DMSO. Formate bound AG1-X2 (50-100 mesh, 250-850μm, Bio-Rad, Hercules, CA) beads of about 250-350μm were washed in DMSO at room temperature for about ten minutes before binding small molecule inhibitors. 1mg/ml recombinant human FGF4 (R&D Systems, Minneapolis, MN) was re-suspended in 0.1% filter sterilized BSA in 1x PBS. Heparin acrylic beads about 250-350 μm (Sigma-Aldrich, St. Louis, MO) were used to deliver FGF4 to duck embryos. A 160μg/ml solution containing equal parts recombinant human TGFβ2 and TGFβ3 (R&D Systems, Minneapolis, MN) was prepared using filter sterilized 4mM HCl in PBS containing 0.1% BSA. Affigel Blue beads about 250-300μm (50-100 mesh, 150-300μm, BioRad, Hercules, CA) were used to deliver TGFβ ligands to quail and duck embryos. Both FGF4 bound heparin acrylic beads and TGFβ2 and TGFβ3 bound Affigel Blue beads were implanted into duck embryos to deliver a combination of all three ligands. Beads were soaked in small molecule inhibitors or ligands for one hour at room temperature before implantation. All concentrations were based on those used previously (Eames and Schneider, 2008; Hayamizu et al., 1991; Niswander et al., 1993; Schneider et al., 2001). Stage HH32 and HH33 embryos were housed in room temperature incubators for one hour before surgeries to minimize embryonic motility. For each bead type used, control surgeries were conducted using beads to deliver carrier. All surgically implanted embryos were collected at HH38. Cleared and stained cases with extensive cartilage and/or bone defects were excluded from analysis under the assumption that a malformation in the jaw skeleton would adversely affect the native mechanical environment. Two-tailed Fisher’s exact test was used to determine statistical significance (Prism 7, GraphPad).

### Endoscopy and jaw motility quantification

*In ovo* video footage of quail and duck from HH32 to HH38 was recorded while eggs incubated at 37.5°C. Video recordings were captured using a 1088 HD High Definition Camera (Stryker, Kalamazoo, MI) with a 4mm, 30° arthroscope (Stryker, Kalamazoo, MI). A universal, dual-quartz, halogen, fiber-optic light source (CUDA Surgical, Jacksonville, FL) was threaded onto the endoscope to provide illumination. The arthroscope was inserted through a small opening in the incubation chamber until it was submerged in albumin. Embryos were acclimated to the light source for 15 minutes prior to recording. Three 10-minute videos were collected from each embryo. The interval of time from the first jaw movement to 5 seconds after the last jaw movement was defined as an activity period, similar to a published quantification method (Hamburger et al., 1965). Average percent active time was calculated along with 95% confidence intervals. Significance was determined using an unpaired, two-tailed Holm-Sidak test adjusted for multiple comparisons (Prism 7, GraphPad).

### 3D reconstruction and finite element analysis

To characterize species-specific differences in the biomechanical environment of the jaw adductor complex, linear finite element analysis (FEA) was used to predict the magnitude and distribution of the von Mises stress on the CP at the adductor insertion. HH33 mandibles from duck and quail were serially sectioned (10μm thickness), stained with Milligan’s trichrome, and imaged at 2.5X magnification. Images were aligned using the orbit and Meckel’s cartilage as landmarks. Meckel’s, the quadrate, surangular, and the MA were manually segmented and reconstructed in 3D (Amira 6; FEI, Hillsboro, OR). The resulting 3D reconstructions of the jaw complexes were imported into commercial FEA software (ANSYS 17; Canonsburg, PA), which was used for meshing and analysis. Tissues were meshed using tetrahedral elements, which were sized based on convergence results from an iterative mesh refinement procedure. Final models utilized 178,378 (duck) and 54,954 elements (quail). The material properties calculated by Tanck et al. (2000) for mineralized embryonic mouse metatarsals (Young’s Modulus (E) = 117MPa; Poisson’s Ratio (v) = 0.3) were used for the surangular and Meckel’s. The other structures were suppressed prior to performing FEA. Boundary conditions were prescribed to mimic those arising during jaw gaping, and included: 1) a fixed support at the contact surface between Meckel’s and the quadrate; and 2) tensile force (duck 3.28E-04 N; quail 1.05E-04 N) aligned with the longitudinal axis of the MA. The magnitudes of the adductor forces were determined using cross-sectional area measurements performed at the longitudinal midpoints and an assumed tensile stress of 1.11kPa (Landmesser and Morris, 1975). Statistical significance was determined using an unpaired, two-tailed, t-test (Prism 7, GraphPad).

### Embryo paralysis

HH32 or HH33 duck were paralyzed using 10mg/ml decamethonium bromide (DMBr) (Sigma-Aldrich, St. Louis, MO) in Hank’s Buffered Sterile Saline (HBSS) and filter sterilized using a 0.22μm filter. Each embryo was treated with a 0.5ml dose of the DMBr solution administered as previously described (Hall, 1986; Solem et al., 2011).

### Microdissections, RNA extraction, RT-qPCR, and analysis

MA insertions were dissected from paralyzed and control duck embryos at HH33 and HH36 and snap frozen in 70% EtOH mixed with dry ice. Microdissected samples were homogenized using a bead-mill (Omni International, Kennesaw, Kentucky) and RNA was isolated using the ARCTURUS PicoPure RNA Isolation Kit (ThermoFisher, Waltham, MA). 200ng cDNA libraries were generated from RNA samples using iScript reverse transcriptase (BioRad, Hercules, CA). Each cDNA library was subsequently diluted to 2ng/μl. Duck *MYOD1, SOX9, TN-C*, and *UCHL-1* primer pairs were used to determine the relative enrichment of muscle, cartilage, tendon, and nerve tissues, respectively, relative to cDNA libraries from duck jaw complexes (Table S1). For quality control, HH33 cDNA libraries were excluded from analysis if the sample was enriched for muscle (>1-fold enrichment of *MYOD1* over control cDNA libraries), nerve (>1.5-fold enrichment of *UCHL-1* over control cDNA libraries), or tendon (>2.5-fold enrichment of *SOX9* over control cDNA libraries). At HH36, the top six tendon enriched samples with less than 4-fold *MYOD1* enrichment were included in the analyses. *Fgf2, Fgf4, Fgf8, Fgfr1, Fgfr2, Fgfr3, Pea3, Erm, Tgfβ2, Tgfβ3, Tgfβr1, Tgfβr2, Tgfβr3, Smad3, Smad7b, and Pai1* expression was quantified by RT-qPCR using duck-specific primer pairs (Table S1). For all genes, expression was normalized to *β-Actin* and analysis was done following the ΔΔC(t) method (Ealba and Schneider, 2013; Livak and Schmittgen, 2001). P-values for - ΔΔC(t) values were calculated using an unpaired, two-tailed, Holm-Sidak test adjusted for multiple comparisons (Prism 7, GraphPad).

### Generation of chimeras

GFP-chick (Crystal Bioscience, Emeryville, CA) and white Pekin duck eggs were incubated to HH9. Tungsten needles and Spemann pipettes were used to graft two differently sized populations of NCM from chick donors into stage-matched duck hosts, producing chimeric “chuck” (Fish and Schneider, 2014a; Fish et al., 2014; Merrill et al., 2008; Schneider, 1999; Schneider and Helms, 2003; Tucker and Lumsden, 2004). Small grafts extended from the middle of the midbrain to the rostral hindbrain at rhombomere 2, whereas large grafts extended from the forebrain–midbrain boundary to rhombomere 2. Comparable-sized regions were excised from duck hosts. Orthotopic grafts and sham operations were performed as controls. Controls and chimeras were incubated side-by-side to ensure accurate staging during collections.

## Results

### Adult jaw morphology is presaged during embryonic development

There are many species-specific differences between Japanese quail and white Pekin duck mandibles. Quail mandibles are slender with a smooth CP and diminutive retroarticular process (Fig.1A). Duck mandibles feature a robust, laterally protruding CP. Furthermore, duck mandibles are larger than quail, both absolutely and in relative proportion, and have a sizeable retroarticular process (Fig.1B). Clearing and staining reveals that species-specific jaw morphology is established during embryonic development (Fig.1C,D). At HH38, an elongate Meckel’s cartilage is surrounded by lower jawbones, and the retroarticular processes are largely comprised of cartilage, yet quail and duck morphologies are already distinguishable. The most obvious difference is a secondary cartilage intermediate within the MA insertion along the surangular in duck. Such cartilage is visible in cleared and stained duck as early as HH36. A secondary cartilage never forms on quail or chick CP.

**Fig.1.**
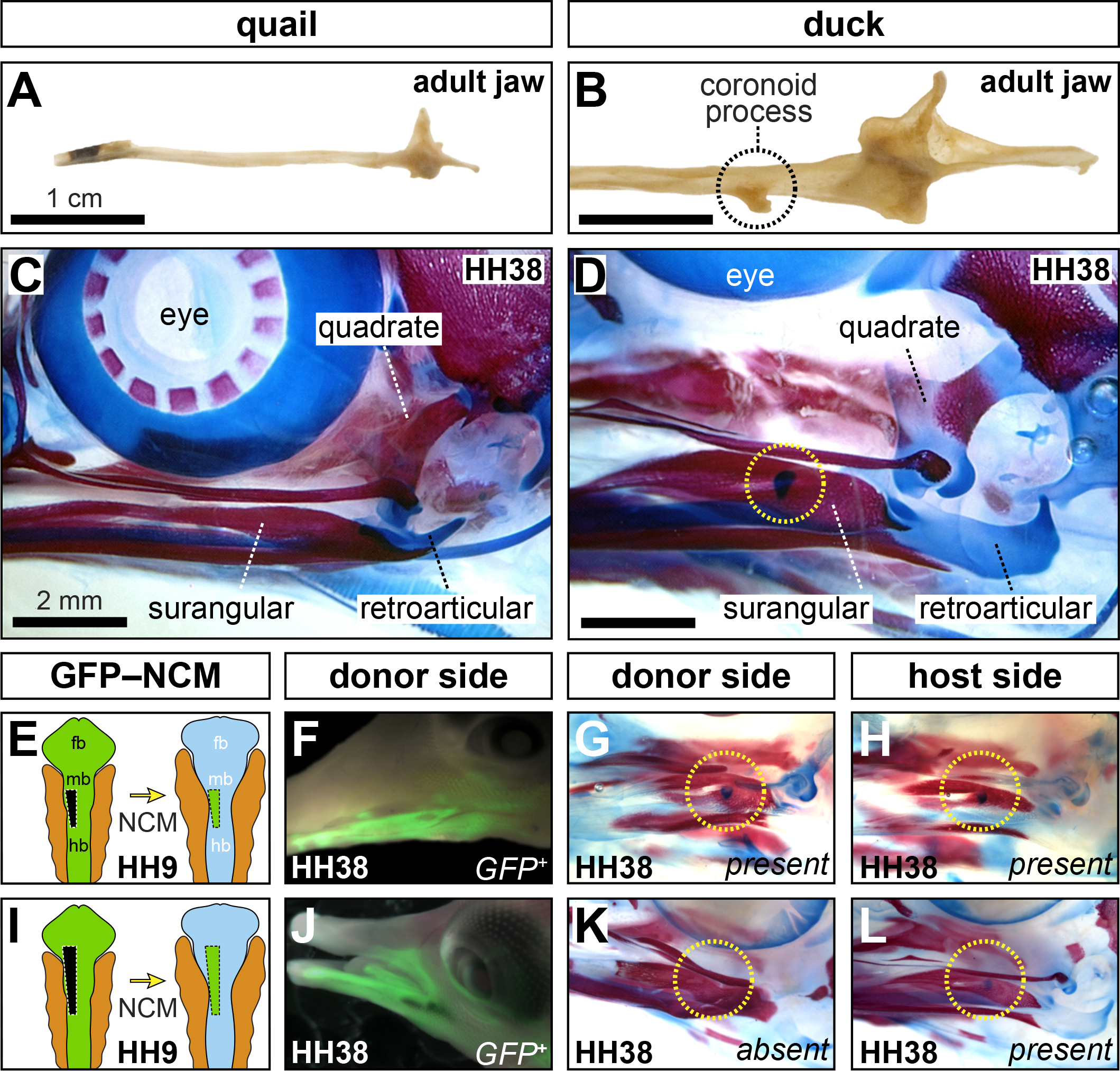
Species-Specific form of the jaw and role of NCM. (**A,B**) Ventral views of left mandibles reveal the smooth appearance in quail and laterally protruding CP in duck (dashed circle). (**C,D**) Left lateral views of cleared and stained skulls showing cartilage (blue) and bone (red). A secondary cartilage forms on the lateral surface of the surangular in duck but not in quail. (**E**) Chimeric “chuck” were produced by unilaterally transplanting small NCM grafts from the midbrain and hindbrain of a GFP-positive chick donor into a comparable position in a stage-matched duck-host. (**F**) Small GFP-chick transplants yield a limited distribution of NCM-derived connective tissues. (**G,H**) The chick-donor side shows little transformation and resembles the contralateral control duck side with secondary cartilage present. (**I,J,K,L**) Larger NCM grafts distribute GFP-positive cells more broadly and lead to a loss of secondary cartilage relative to the contralateral, duck-host side.

### NCM patterns the MA complex in a dose-dependent manner

NCM transplanted from HH9 GFP-positive chick into stage-matched duck hosts transforms the morphology of the jaw and CP (Fig.1E,F,I,J). The extent of transformation and distribution of GFP-positive NCM-derived connective tissues depends upon donor graft size. Small NCM transplants result in a limited distribution of GFP-positive skeletal and connective tissues, and produce minor changes to the size and shape of the jaw skeleton, but not enough to affect the secondary chondrogenesis (Fig.1G,H). In contrast, large transplants result in extensively distributed GFP-positive skeletal and connective tissues, and transform the jaw to become more chick-like, including the absence of a secondary cartilage on the donor side CP (Fig.1K,L).

### The progression of embryonic jaw motility is similar in quail and duck

*In ovo* videos of embryonic jaw motility captured periodic jaw gaping in quail and duck embryos (Fig.2A,B,C,D)(Movie S1,S2). The first quantifiable jaw movements occur at HH33 in quail and duck. HH33 quail are active 10.46% of the time (95% CI ±3.07%, n=9) while stage-matched duck are active 5.2% of the time (95% CI ±1.06%, n=10). Both the frequency and duration of jaw movements increase with developmental time in quail and duck (Fig.2E,F). Quail and duck jaw motility track closely at HH34 (18.82%±8.32%, n=12 for quail and 15.72%±3.28%, n=18 for duck) and HH35 (28.58%±16.63%, n=6 for quail and 29.35%±6.57%, n=2 for duck). No statistically significant differences in motility are observed in developmental stages preceding the appearance of secondary cartilage. A significant difference is observed at HH36 (26.66%±8.36%, n=22 for quail, and 43.97%±5.06, n=26 for duck, p<0.0005), however, by this stage, a secondary cartilage is already formed on the CP. Peak quail jaw motility is observed at HH37 (67.39%±5.7%, n=6 in quail, versus 51.72%±8.69%, n=13 in duck) while duck motility peaks at HH38, but does not exceed quail motility (60.76%±5.79%, n=7 in duck versus 61.67%±5.49%, n=7 in quail).

**Fig.2.**
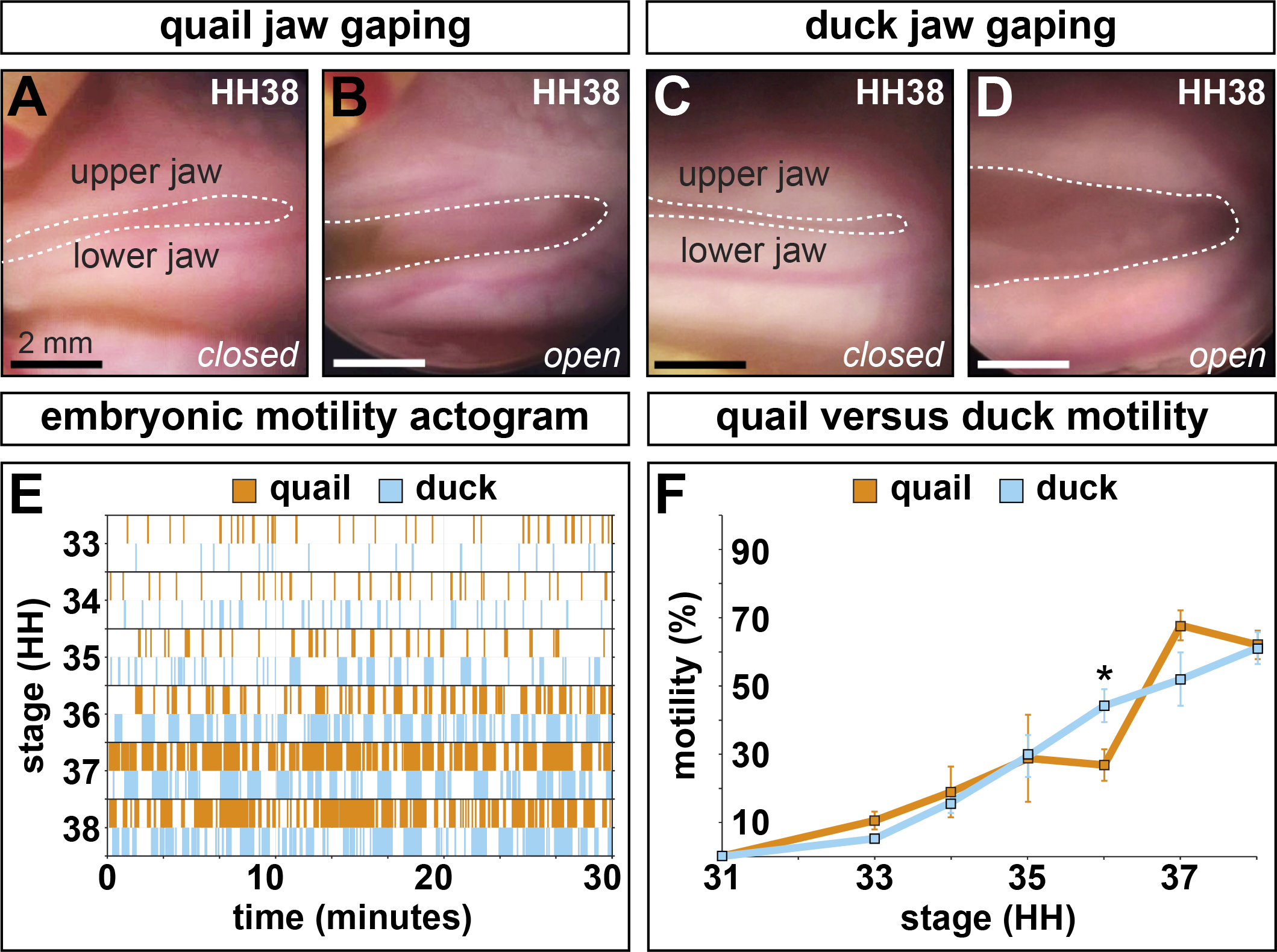
Jaw motility *in ovo.* (**A**,**B**,**C**,**D**) Representative open and closed jaw gaping positions in quail and duck embryos. (**E**) Actogram of 30-minute observation periods for representative quail and duck. Six consecutive stages were observed. Quail and duck activity periods steadily increase in frequency and duration. (**F**) During HH33, a key stage of secondary cartilage induction, the differences in jaw motility are minimal with quail being slightly more active, though the difference is not significant. Duck are significantly more active at HH36 (p<0.0005).

### FEA predicts distinct mechanical environments at the quail and duck coronoid process

3D reconstructions of HH33 quail and duck jaws including Meckel’s, the quadrate, postorbital, surangular, and MA were created by manually segmenting histological images (Fig.3A,B). Reconstructions reveal species-specific, geometrical differences in cross-sectional area of the muscle, direction of contractile force, and area of the surangular over which force is applied. In duck, the MA inserts on the lateral aspect of the surangular, while in quail, the insertion is dorsal. In duck, the insertion is also more proximal to the jaw joint. At its widest, the cross-sectional area of the duck MA is 321,000μm^2^, while the slender quail muscle is only 114,192μm^2^ indicating the maximum contractile force of the duck muscle is roughly 2.8 times greater than in quail.

**Fig.3.**
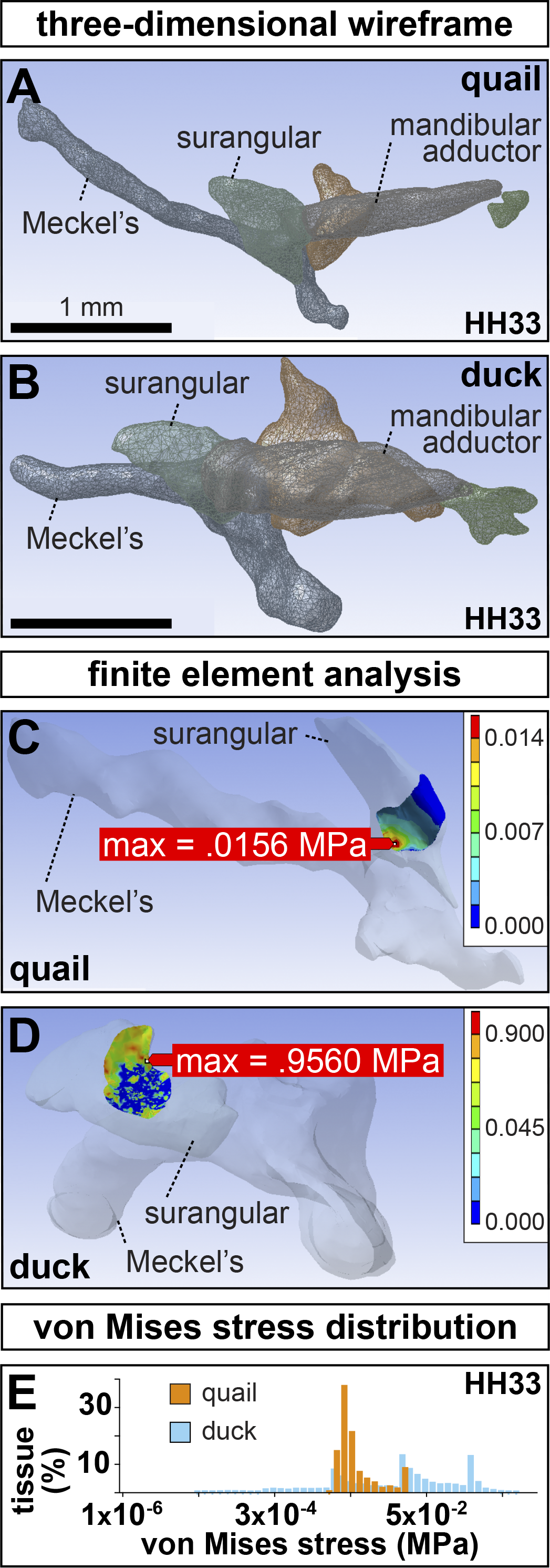
3D reconstructions and finite element analysis of the adductor complex. Three-dimensional wireframes of left (**A**) quail and (**B**) duck jaw showing the presumptive surangular (light-green), quadrate (red), MA muscle (purple), post-orbital (dark-green), and Meckel’s (blue). Note the slender MA and its dorsal insertion on the quail surangular versus the bulky MA and its lateral insertion in duck. (**C**) Finite element modeling predicts a maximum von Mises stress concentration of 0.0156 MPa within the medial portion of the contact area between the MA and the surangular in quail. Color scales indicate predicted von Mises stress. (**D**) A maximum von Mises stress concentration of 0.9560 MPa is predicted within a dorsolateral region in duck. (**E**) Histogram of the range of von Mises stresses in duck versus quail. Note that the maximum von Mises stress in quail is substantially less than in duck.

Finite element models of the insertion site between the MA and the surangular predict that duck experience a maximum shear stress concentration roughly 60 times greater than quail (0.96MPa in duck versus 0.016MPa in quail)(Fig.3C,D). Furthermore, the mean von Mises stress experienced in duck (0.053MPa) is significantly higher than in quail (0.0045MPa; p<0.0001). Histograms also reveal the state of shear stress at the insertion is more homogeneous in quail, while tissue at the duck insertion is subjected to a broader range of shear stress (Fig.3E).

### The FGF pathway changes during development and is affected by paralysis

RT-qPCR analyses on microdissected duck MA insertions reveal significant increases in ligands *Fgf2* (5.34±1.50-fold change, p<0.0005), *Fgf4* (449.89±237.59-fold change, p<0.0005), and *Fgf8* (56.22±44.55-fold change, p<0.0005) from HH33 to HH36 (n=13 for HH33 controls, n=10 for HH36 controls)(Fig.4A). FGF receptors *Fgfr1* (0.76±0.21-fold change, p<0.05), *Fgfr2* (0.19±0.18-fold change, p<0.0005), and *Fgfr3* (0.68±0.30-fold change, p<0.05) significantly diminish in expression over this time. Transcriptional effectors *Pea3* (5.61±1.09-fold change, p<0.0005) and *Erm* (2.44±0.54-fold change, p<0.0005) are both significantly more abundant at HH36 than at HH33.

**Fig.4.**
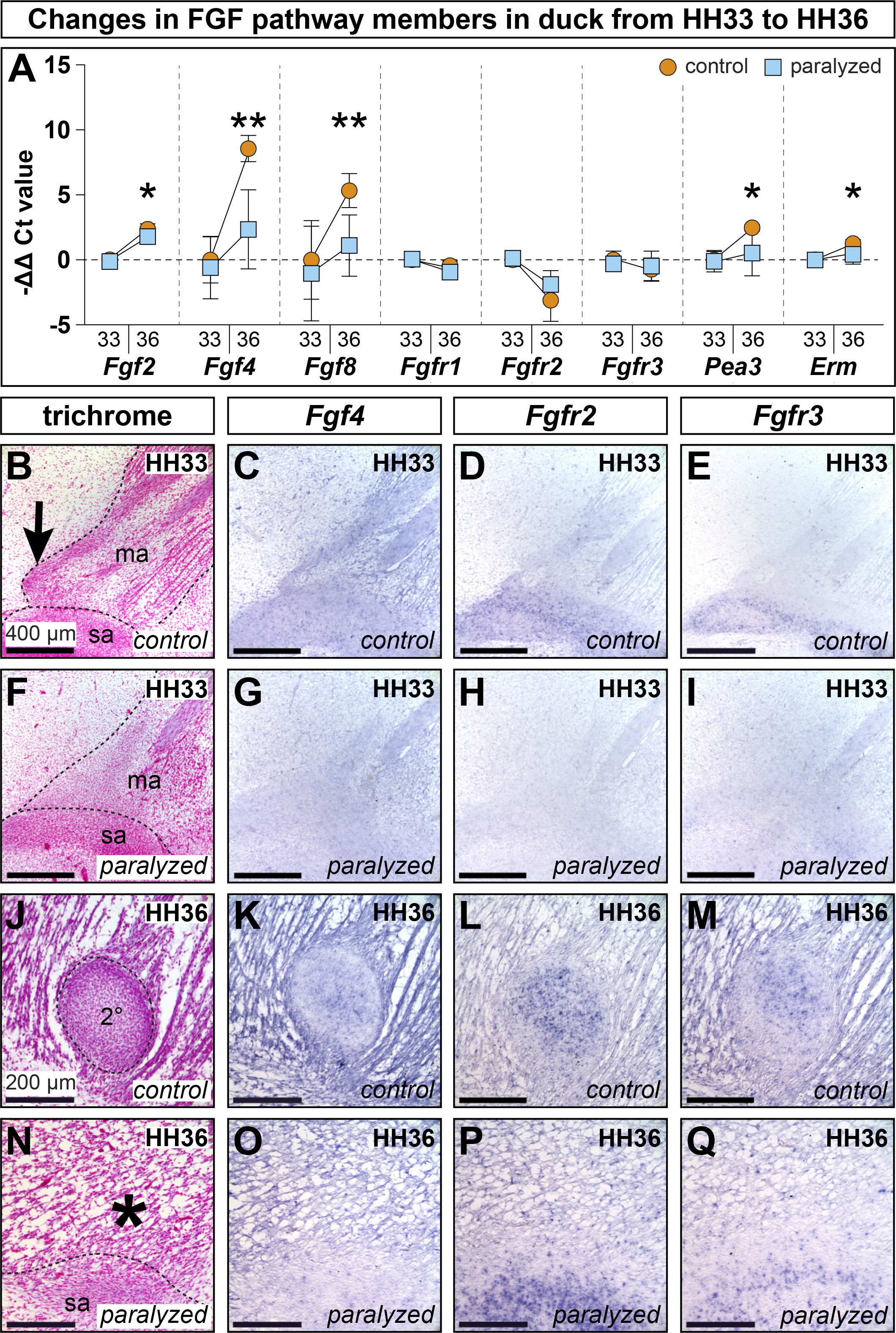
FGF pathway in paralyzed and control duck. (**A**) Differential expression in isolated MA entheses from HH33 and HH36 control and paralyzed embryos. Each gene is normalized to *β-Actin* and shown relative to HH33 controls. Error bars represent standard deviation. Asterisks denote statistical significance between control and paralyzed samples at HH36 (*p<0.05; **p<0.005). (**B**) Sagittal section through the MA (ma) muscle insertion along the presumptive surangular (sa). A secondary cartilage condensation is present at the MA insertion on the CP (arrow). (**C,D**) *Fgf4* and *Fgfr2* (stained purple) are expressed in the secondary cartilage condensation and surrounding tissues. (**E**) *Fgfr3* is expressed around the margins of the surangular condensation. (**F**) 24 hours after paralysis at HH32, HH33 embryos show disrupted muscle and tendon, and there is no secondary cartilage condensation. (**G,H**) *Fgf4* and *Fgfr2* are altered and the secondary cartilage is absent. (**I**) *Fgfr3* is disrupted. (**J**) Sagittal section through the MA muscle insertion on the CP lateral to the surangular. The secondary cartilage (2°) is well formed. (**K,L,M**) *Fgf4, Fgfr2*, and *Fgfr3* are in the secondary cartilage and surrounding tissues. (**N**) Paralysis at HH32 prevents secondary cartilage formation (asterisk). The MA inserts directly onto the surangular. (**O,P,Q**) *Fgf4, Fgfr2, and Fgfr3* are altered and secondary cartilage is absent.

Paralysis at HH32 does not result in significant changes to FGF signaling pathway members or effectors at HH33 relative to stage-matched controls. In HH36 paralyzed embryos, the only FGF ligand with a significant increase is *Fgf2* relative to HH33 controls (3.67±1.30-fold change, p<0.0005)(n=12 for HH33 paralyzed, n=11 for HH36 paralyzed). However, *Fgf2* at HH36 is still significantly less in paralyzed embryos than in stage-matched controls (p<0.05)(asterisk, Fig.4A). In paralyzed HH36 embryos, *Fgf4* is 21.49±33.68-fold more abundant than in HH33 controls and *Fgf8* is 4.79±5.06-fold more abundant, but both genes are still significantly less expressed than in stage-matched controls (p<0.005 for both)(asterisks, Fig.4A). At HH36, *Fgfr1* (0.55±0.22-fold change, p<0.0005) and *Fgfr2* (0.35±0.29-fold change, p<0.0005) are significantly down in paralyzed samples, similar to expression dynamics seen in controls over the same period. Unlike control samples, *Pea3* (2.58±2.75-fold change) and *Erm* (1.49±0.67-fold change) remain relatively flat in paralyzed embryos and, by HH36, are significantly less abundant than in HH36 controls (p<0.05 for both)(asterisks, Fig.4A).

Analysis of spatial and temporal gene expression patterns was conducted in control and paralyzed duck at HH33 and HH36 (Table 1). At HH33, in sagittal section, the MA is visible as two muscle bundles divided proximodistally by the mandibular branch of the trigeminal nerve (Fig.4B). Proximal to the mandibular nerve, the MA appears fan-like and inserts broadly. Distal to the nerve, unipinnate muscle fibers are joined by a fibrous aponeurosis. The musculature and aponeurosis appear relatively disorganized following 24 hours of paralysis (Fig.4F).

**Table 1.**
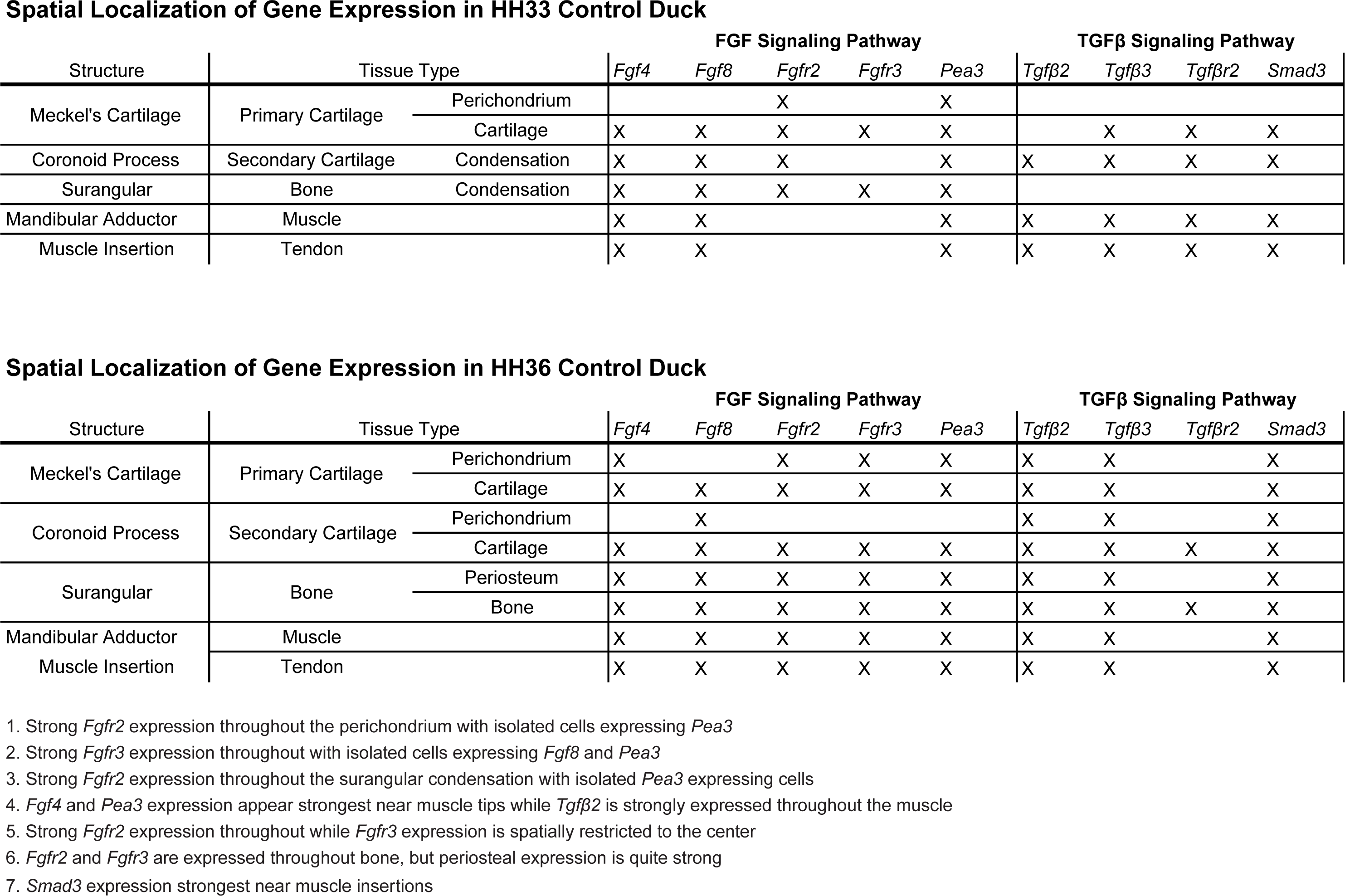

At HH33, *Fgf4* is expressed throughout primary cartilages like the quadrate, and Meckel’s, as well as in skeletal muscles like the MA, the MA insertion, and the mesenchymal condensation that will give rise to secondary cartilage (n=5 for each gene)(Fig.4C). After 24 hours of paralysis, *Fgf4* is maintained in the quadrate and Meckel’s, but diminished in the MA and its insertion (Fig.4G). *Fgf8* is in the MA, the MA insertion, the secondary cartilage insertion, and the surangular condensation (Fig.S1). There is also *Fgf8* in primary cartilages like Meckel’s and the quadrate. The secondary cartilage condensation and its *Fgf8* domain are not present in embryos 24 hours after paralysis (Fig.S1). *Fgfr2* is in the quadrate and Meckel’s, particularly in the perichondrium, as well as in the secondary cartilage condensation and the nascent surangular (Fig.4D). Following 24 hours of paralysis, expression in primary cartilage is maintained, while expression in the secondary cartilage condensation and surangular condensation are diminished (Fig.4H). *Fgfr3* is in the quadrate and Meckel’s, but not perichondria, and in the surangular condensation with greater expression around the periphery (Fig.4E). Paralysis leads to decreased expression in the surangular condensation while expression in primary cartilage is maintained (Fig.4I). *Pea3* is in the MA, the MA insertion and the secondary cartilage condensation (Fig.S1). There is also expression in the surangular condensation, primary cartilages and perichondria. 24 hours after paralysis, the secondary cartilage condensation fails to form and the corresponding region of *Pea3* is absent (Fig.S1).

By HH36, secondary cartilage is present within the MA insertion and is encapsulated in a dense fibrous sheath (Fig.4J). The MA muscles have begun to separate into distinct superficial sheet-like, proximal fan-like, and distal groups of fibers. HH36 paralyzed embryos have poor muscle and tendon organization and lack a secondary cartilage condensation (Fig.4N). *Fgf4* (n=5 for each gene) is strongly expressed at HH36 in the MA, the MA insertion, and the surangular and periostea (Fig.4K). The quadrate and Meckel’s also express *Fgf4* throughout the cartilage and perichondrium. *Fgf4* is also seen within the secondary cartilage condensation. Paralysis prevents secondary chondrogenesis, however, *Fgf4* is maintained in muscle, bone, and primary cartilages (Fig.4O). *Fgf8* is in the MA, tendon, and secondary cartilage (Fig.S1). *Fgf8* is also in the surangular, periosteum, and primary cartilage. Paralysis prevents secondary cartilage from forming, but *Fgf8* is still in muscle and its connective tissues (Fig.S1). *Fgfr2* is in muscle, tendon, bone, periostea, cartilage, perichondria, and within secondary cartilage (Fig.4L). Following paralysis, the only change to *Fgfr2* is the absence of a secondary cartilage domain (Fig.4P). *Fgfr3* is in the quadrate and Meckel’s as well as in the periosteum of the surangular. *Fgfr3* is also in muscle, tendon, bone, periostea, cartilage, perichondria, and secondary cartilage (Fig.4M). Expression in the secondary cartilage is highest at the center and grows lower towards the periphery. In paralyzed embryos, only the *Fgfr3* domain in secondary cartilage is absent (Fig.4Q). *Pea3* is in the MA muscle, tendon, and the secondary cartilage condensation (Fig.S1). *Pea3* is also in primary cartilage, perichondria, bone, and periostea. As secondary cartilage fails to form in HH36 paralyzed embryos, *Pea3* is absent (Fig.S1).

### The TGF*β* pathway changes during development and is affected by paralysis

Quantitative RT-PCR shows that *Tgfβ2* (4.28±1.29-fold change, p<0.0005) and *Tgfβ3* (7.19±2.11-fold change, p<0.0005) increase significantly from HH33 to HH36 (n=10 for HH33 controls, n=10 for HH36 controls)(Fig.5A). Paralyzed embryos mirror the increases in *Tgfβ2* (2.87±1.36-fold change, p<0.05) and *Tgfβ3* (5.50±2.30-fold change, p<0.0005) over the same period. Transcriptional activity of receptors *Tgfβr1, Tgfβr2, Tgfβr3*, and transcriptional effectors *Smad3, Smad7b*, and *Pai1* remain flat in controls. In contrast, HH36 paralyzed samples express more *Pai1* (2.53±1.89-fold change) than HH33 controls (p<0.05), and achieve significantly greater expression than HH36 control samples (p<0.05)(asterisk, Fig.5A).

**Fig.5.**
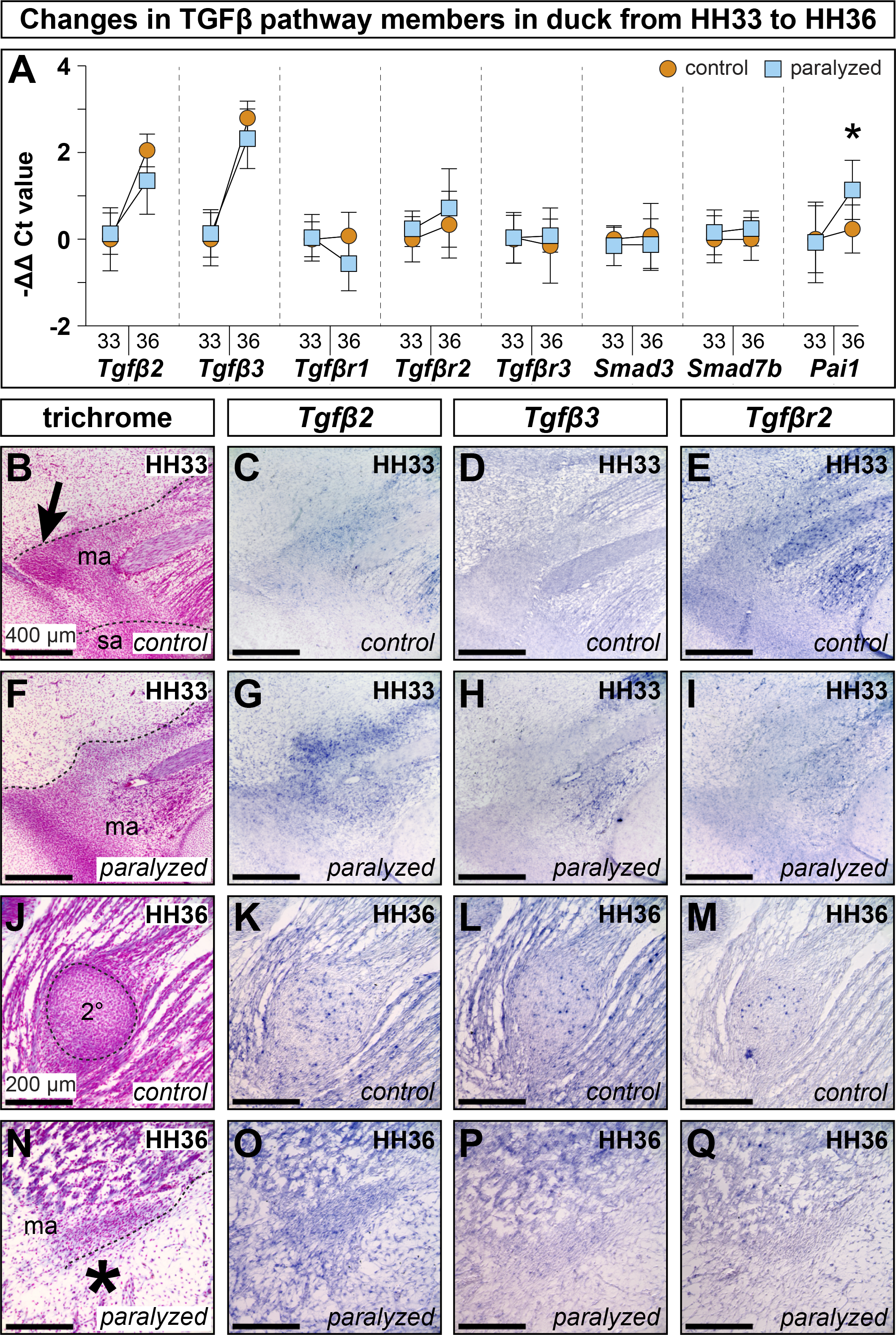
TGFβ pathway in paralyzed and control duck. (**A**) Differential expression in isolated MA entheses from HH33 and HH36 control and paralyzed embryos. Each gene is normalized to *β-Actin* and displayed relative to HH33 controls. Error bars represent standard deviation. Asterisk denote statistical significance between control and paralyzed samples at HH36 (*p<0.05). (**B**) Sagittal section through the MA (ma) muscle insertion along the presumptive surangular (sa). A secondary cartilage condensation is present at the MA insertion on the CP (arrow). (**C,D,E**) *Tgfβ2, Tgfβ3*, and *Tgfβr2* are expressed in the secondary cartilage condensation and surrounding tissues. (**F**) 24 hours after paralysis at HH32, HH33 embryos show disrupted muscle and tendon, and there is no secondary cartilage condensation. (**G,H,I**) *Tgfβ2, Tgfβ3*, and *Tgfβr2* are disrupted. There is no secondary cartilage condensation. (**J**) Sagittal section through the MA muscle insertion on the CP lateral to the surangular. The secondary cartilage (2°) is well formed. (**K,L,M**) *Tgfβ2, Tgfβ3*, and *Tgfβr2* are expressed in the secondary cartilage and surrounding tissues. (**N**) Paralysis at HH32 prevents secondary cartilage formation (asterisk). (**O,P,Q**) *Tgfβ2, Tgfβ3*, and *Tgfβr2* are altered and secondary cartilage is absent.

Our qualitative analyses show that at HH33, *Tgfβ2* is expressed in the MA muscle, the MA insertion, and the secondary cartilage condensation (Fig.5B,C). At HH33, following 24 hours of paralysis, expression in muscle and tendon persists while the secondary cartilage condensation and its *Tgfβ2* domain does not (Fig.5F,G). *Tgfβ3* is also in the MA muscle, the MA insertion, primary cartilage like Meckel’s and the quadrate, and the secondary cartilage condensation (Fig.5D). At this stage, the only *Tgfβ3* domain affected by paralysis is in the secondary cartilage condensation (Fig.5H). *Tgfβr2* is in the MA, the MA insertion, and in the secondary cartilage condensation (Fig.5E). *Tgfβr2* is also in Meckel’s and the quadrate. Following paralysis, the only expression domain affected is the secondary cartilage condensation (Fig.5I). *Smad3* is in the MA, the insertion, and the secondary cartilage condensation (Fig.S1). *Smad3* is also in the quadrate, Meckel’s, and other primary cartilages. The secondary cartilage domain does not appear in stage-matched, paralyzed embryos (Fig.S1).

In HH36 duck, *Tgfβ2* is in muscles like the MA, tendons like the MA insertion, bones like the surangular and their periostea, and cartilages like Meckel’s, the quadrate, and their perichondria (Fig.5K). *Tgfβ2* is also expressed throughout the secondary cartilage on the CP. Following paralysis, the only change in expression at HH36 is for *Tgfβ2* coincident with the loss of secondary cartilage (Fig.5O). *Tgfβ3* is in all the same tissues as *Tgfβ2* in HH36 control and paralyzed embryos, including the secondary cartilage (Fig.5L,P). By HH36, *Tgfβr2* is in the surangular, as well as secondary cartilage on the CP (Fig.5M). Following paralysis, the secondary cartilage and its *Tgfβr2* domain are absent while *Tgfβr2* in bone is unaffected (Fig.5Q). *Smad3* is in the MA and its insertion, and in the secondary cartilage. There is also *Smad3* in primary cartilages, perichondria, bone, and periostea (Fig.S1). Paralyzed HH36 embryos do not form secondary cartilage so the corresponding *Smad3* expression is absent (Fig.S1).

### Inhibiting FGF or TGFβ signaling affects the condensation of secondary cartilage

Unilateral delivery of FGF signaling inhibitor SU5402 blocks the formation of, or reduces the size of secondary cartilage on the CP (n=18 at HH32, n=29 at HH33)(Fig.6A,C). No change in secondary cartilage is observed following delivery of DMSO control beads (n=6). The efficacy of secondary cartilage inhibition at HH38 depends upon the stage of treatment, with HH32 embryos being more sensitive to FGF inhibition than HH33 embryos (Fisher’s exact test, p=0.0047). In 88.9% of embryos treated with SU5402 at HH32, secondary cartilage is either lost or reduced in size (n=16/18). Of those secondary cartilage phenotypes, 50% are reduced in size (n=8/16), and 50% have a complete absence (n=8/16) of secondary cartilage. FGF inhibition at HH33 reduces the size of the secondary cartilage in 31.01% of cases (n=9/29) and prevents secondary cartilage induction in 13.79% of cases (n=4/29).

**Fig.6.**
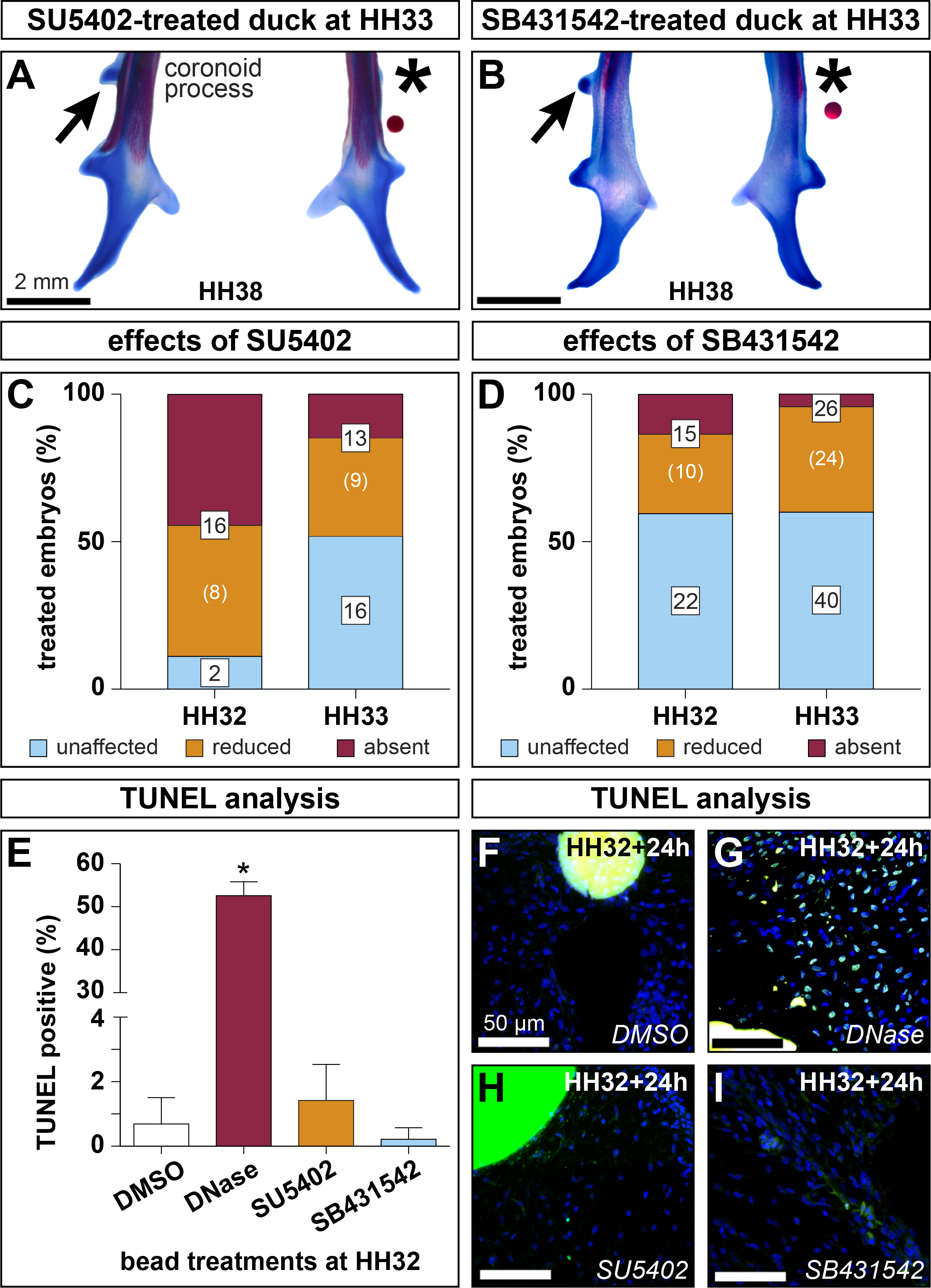
Inhibition of FGF and TGFβ signaling during secondary chondrogenesis. (**A**) Ventral view of a cleared and stained duck mandible treated with a bead soaked in an FGF inhibitor (SU5402). Note the loss of secondary cartilage (asterisk) while the untreated side develops normally (arrow). (**B**) Inhibition of TGFβ signaling (SB431542) results in a loss of secondary cartilage while the control side develops normally. (**C**) FGF signaling inhibition eliminates or reduces secondary cartilage by HH38, with a greater treatment effect at HH32 versus HH33 (Fisher’s Exact Test p<0.005). (**D**) TGFβ signaling inhibition eliminates or reduces secondary cartilage by HH38. (**E**) Inhibiting FGF or TGFβ signaling does not increase apoptosis after 24 hours. Positive control, DNase digested slides displayed significant apoptosis (unpaired t-test p<0.0001). (**F,G,H,I**) Sections from DMSO, SU5402, or SB431542 treated embryos reveal little apoptosis. Extensive positive staining was observed in DNase digested sections.

Inhibition of TGFβ signaling by delivering SB431542 also frequently causes loss or reduction in the size of the secondary cartilage on the CP (n=37 at HH32, n=66 at HH33)(Fig.6 B,D). Although the statistical distribution of outcomes does not depend on whether embryos are treated at HH32 (40.54% absent or reduced secondary cartilage, n=15/37) or HH33 (39.39% absent or reduced secondary cartilage, n=26/66), HH32 treatments tend to be more efficacious at preventing secondary chondrogenesis (13.51%, n=5/37) than HH33 treatments (3.03%, n=2/66).

### Inhibiting FGF or TGFβ signaling does not lead to increased cell death

TUNEL staining shows that implanting AG1X2 chromatography beads soaked in DMSO (n=3 embryos) or small molecule inhibitors of FGF signaling (n=6 embryos) or TGFβ signaling (n=7 embryos) at HH32 does not increase cell death nor did we observe histological evidence at any stage where muscle or tendon formation were blocked by treatment delivery (data not shown). 24 hours after implantation, 0.69% of cells surrounding DMSO soaked beads are undergoing apoptosis (n=5 sections)(Fig.6E,F). There is no significant increase in cell death over control beads with SU5402 (1.42%, n=19 sections) or SB431542 (0.22%, n=29 sections)(Fig.6H,I) treatments. For comparison, DNase-treated positive control slides show significantly more cell death (52.60%, n=3 sections, unpaired t-test p<0.0001)(Fig.6G).

### Exogenous FGF and TGFβ treatments can restore cartilage in paralyzed embryos

HH38 duck embryos paralyzed and treated with FGF4 beads at HH32 form cartilage adjacent to or surrounding the bead in 27.27% of cases (n=3/11)(Fig.7B). No cartilage is induced in any embryos treated with BSA beads alone (n=4 heparin acrylic, n=12 Affigel blue)(asterisk, Fig.7A), or in cases where recombinant protein soaked beads are located far from the MA insertion (n=4 for FGF4, n=2 for TGFβ2/TGFβ3, and n=4 for FGF4/TGFβ2/TGFβ3). Paralysis and implantation of beads soaked in TGFβ2 and TGFβ3 induce cartilage in 75% of HH38 duck (n=15/20)(Fig.7C). Implanting both FGF4 and TGFβ2/TGFβ3 soaked beads in paralyzed HH32 duck induces cartilage in 85.71% of cases (n=12/14)(Fig.7D). Treating HH32 quail with exogenous TGFβ2/TGFβ3 induces a chondrogenic response in 11.11% of embryos (n=1/9)(Fig.7E). Safranin-O staining confirms the presence of a glycosaminoglycan-rich cartilaginous extracellular-matrix surrounding the beads (n=2/3)(Fig.7F). Although spherical beads were implanted, the axial orientation of Safranin-O-positive tissue surrounding the beads is not radially symmetrical and tends to align with the orientation of the MA insertion. Analysis of paralyzed duck rescue experiments reveal that the distribution of phenotypes depends upon the ligand or ligands received (Fisher’s Exact Test, p=0.005)(Fig.7G).

**Fig.7.**
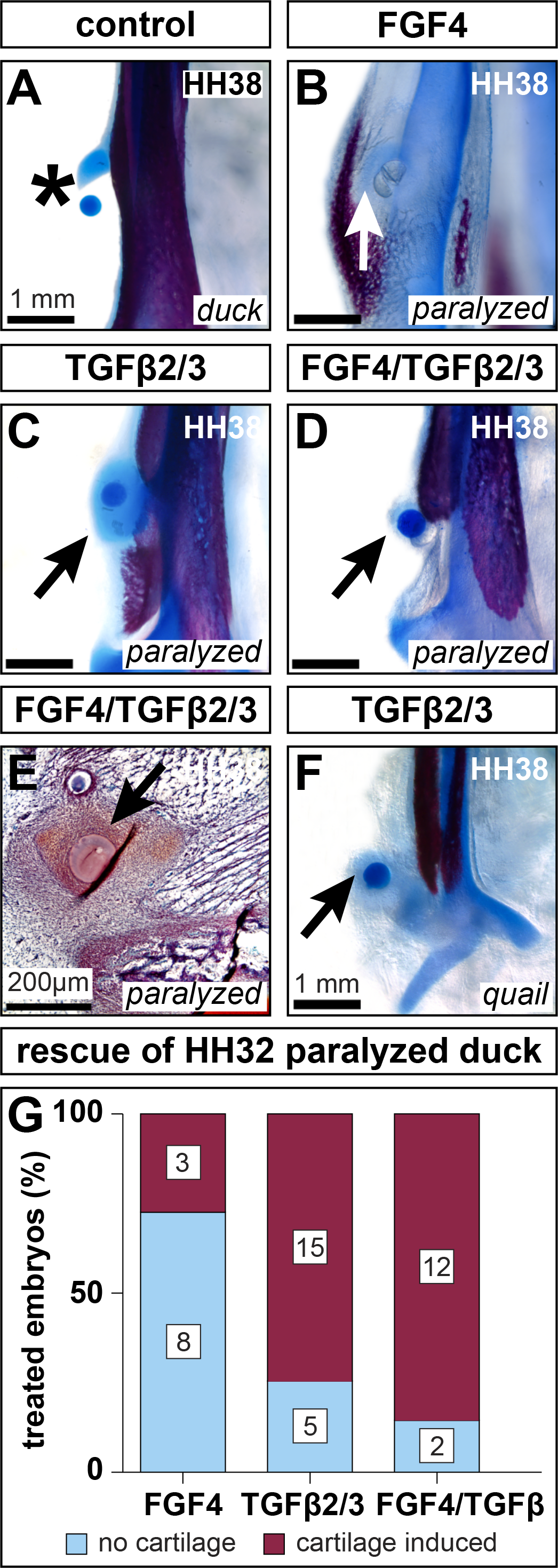
FGF4 and TGFβ2/TGFβ3 induce chondrogenesis. (**A**) Ventral view of a cleared and stained mandible treated with a BSA soaked bead. Carrier treatments exert no effect on secondary cartilage (asterisk). (**B**) HH32 FGF4 treatment induces cartilage (arrow) in paralyzed embryos by HH38. (**C**) TGFβ2/TGFβ3 treatment induces cartilage (arrow) in paralyzed embryos. (**D**) Combined FGF4 and TGFβ2/TGFβ3 treatments induce cartilage (arrow) despite paralysis. (**E**) HH38 sagittal section through the MA insertion of a paralyzed embryo implanted with FGF4 and TGFβ2/TGFβ3 beads at HH32. Safranin-O reveals dense, positively stained mesenchyme surrounding the beads (arrow). (**F**) HH32 TGFβ2/TGFβ3 treatment induces quail to form cartilage by HH38 (arrow). (**G**) FGF4, TGFβ2/TGFβ3, and FGF4/TGFβ2/TGFβ3 treatments induce cartilage by HH38. The distribution of treatment outcomes depends upon the ligand or ligands embryos receive (Fisher’s Exact Test p=0.005).

## Discussion

### NCM controls the species-specific pattern of the MA insertion

In previous studies we have shown that NCM controls the species-specific size and shape of the jaw skeleton and associated musculature via cell-autonomous morphogenetic programs (Solem et al., 2011; Tokita and Schneider, 2009). But in the present study we go further and substantiate that this patterning ability is dose-dependent. While we know that the extent of gene expression in chimeras is directly related to the degree of chimerism (Ealba and Schneider, 2013), here we were able to extend this principle to morphology and modulate the presence or absence of secondary cartilage on the CP by titrating the size of donor NCM transplants and thus the distribution of NCM-derived connective tissues. Small transplants did not alter secondary cartilage development whereas larger transplants did. Based on our prior analyses of muscle and connective tissue patterning (Solem et al., 2011; Tokita and Schneider, 2009), and the critical role for interactions between NCM and muscle precursors (Bothe et al., 2007; Evans and Noden, 2006; Grenier et al., 2009; Noden, 1983, 1988; Noden and Trainor, 2005; Rinon et al., 2007), we expect that increasingly larger populations of donor NCM relocate the MA insertion from a duck-like lateral position to one that is more dorsal and chick-like. In this way, and concomitant with its patterning abilities, NCM would be acting as a major determinant of the mechanical environment whereby specific loading conditions are more conducive to secondary cartilage formation.

### Quality not quantity of mechanical stimulation drives secondary chondrogenesis

Secondary cartilage development can be divided into two phases: induction and maintenance. Both phases require proper biomechanical stimulation. Embryonic motility is an essential source of biomechanical stimulation and the developmentally plastic response to biomechanical loading is a potent mechanism through which embryonic form comes to presage adult function (Anthwal et al., 2015; Blitz et al., 2009; Brunt et al., 2017; Carter and Beaupré, 2007; Hall, 1967, 1968, 1972, 1986; Hall and Herring, 1990; Havis et al., 2016; Huang et al., 2013; Kardon, 1998; Pitsillides, 2006; Pollard et al., 2014; Schweitzer et al., 2010; Sharir et al., 2011; Shwartz et al., 2012; Solem et al., 2011; Wu et al., 2001). For induction of secondary cartilage to occur, the frequency of mechanical stimulation must cross a threshold (Hall, 1967, 1968). The size of a secondary cartilage can also be decreased by paralysis after secondary cartilage induction (Solem et al., 2011). The similarity in early quail and duck jaw motility indicates that frequency of jaw activity is an unlikely determinant of species-specific secondary chondrogenesis. A significant difference in motility manifests at HH36, though a secondary cartilage is already formed in duck by that time. Thus, we conclude that the frequency of mechanical stimulation is not, itself, sufficient to induce secondary cartilage in quail versus duck, which points to the role of biomechanical stress resulting from a combination of species-specific muscle pattern and resultant differences in the quality or type of functional loading on the muscle insertion.

### Mechanical cues result from and contribute to species-specific morphology

Prior work has highlighted the contribution of the mechanical environment in wrap-around and other force-transmitting tendons (Benjamin and Ralphs, 1998; Blitz et al., 2013; Carter and Beaupré, 2007; Murchison et al., 2007; Schweitzer et al., 2010; Shwartz et al., 2013). Such a configuration, in which a tendon experiences not only axial tension, but also compression in which the tendon is held taught against the bone, is conducive to fibrocartilage development (Blitz et al., 2009; Koo et al., 2017). Thus, the evolutionary presence or absence of secondary cartilage on the CP reflects species-specific variation in functional anatomy determined by *in ovo* mechanical loading (Beresford, 1981; Fang and Hall, 1997; Hall, 1979; Stutzmann and Petrovic, 1975). In taxa such as humans, rats, cats, and duck, secondary cartilage forms at the jaw adductor muscle insertion (Amorim et al., 2010; Amorim et al., 2008; Hall, 2005; Horowitz and Shapiro, 1951; Kantomaa and Rönning, 1997; Moore, 1973, 1981; Solem et al., 2011; Soni and Malloy, 1974; Vinkka, 1982; Washburn, 1947) whereas an equivalent secondary cartilage is absent in mice, guinea pigs, chick, and quail (Anthwal et al., 2008; Anthwal et al., 2015; Boyd et al., 1967; Moss and Meehan, 1970; Rot-Nikcevic et al., 2007; Shibata et al., 2003; Solem et al., 2011). Our work implies that the reason secondary cartilage forms at this location in some species and not others is due to the way NCM-mediated muscle pattern leads to differential forces during embryonic motility.

To our knowledge, this is the first finite element modelling of the embryonic jaw adductor complex. Our FEA illuminates the difference in both the predicted magnitude and spatial distribution of von Mises stress in the MA insertion of embryonic quail and duck prior to secondary chondrogenesis. Perhaps the wide ranging magnitudes of shear stress distributed across the surface of the duck surangular mediates the precise biomechanical cues required to elicit a spatially restricted domain of secondary cartilage. The secondary cartilage is the future site of an ossification center that fuses to the surangular, enables robust osteointegration, and further distinguishes both the form and the functional mechanics of the duck versus quail jaw apparatus. However, the mechanisms that facilitate the relationship between mechanical stimulation and musculoskeletal adaptation have remained largely unknown. While previous studies have implicated FGF and TGFβ signaling in both early, muscle-independent, and late, muscle-dependent, phases of sclerotome-derived limb tendons (Havis et al., 2016; Havis et al., 2014; Huang et al., 2015), our findings suggest that mechanical cues drive differential activation of FGF and TGFβ signaling to induce species-specific secondary cartilage within an NCM-derived tendon insertion. Moreover, we do not observe any evidence for crosstalk between these pathways, given that paralysis downregulates FGF signaling while TGFβ expression remains unchanged. Conversely, despite the maintenance of TGFβ, FGF is downregulated. Such findings are consistent with the independent functions of these pathways during chick limb tendon morphogenesis (Havis et al., 2016). However, manipulating these pathways in the limb has not been shown to induce cartilage formation.

### FGF and TGFβ are necessary and sufficient for secondary chondrogenesis

Molecular programs of tendon development are context-dependent. In mouse limbs, TGFβ signaling promotes tendon development while FGF signaling is inhibitory (Blitz et al., 2013; Havis et al., 2014; Pryce et al., 2009; Subramanian and Schilling, 2015). However, FGF signaling is a pro-tendon signal in chick limbs and promotes axial mouse and chick tendon development (Brent et al., 2005; Brent et al., 2003; Edom-Vovard et al., 2001; Edom-Vovard et al., 2002; Havis et al., 2016; Havis et al., 2014; Smith et al., 2005). Our quantitative and qualitative analyses demonstrate that FGF and TGFβ ligands, receptors, and effectors are expressed in musculoskeletal tissues throughout stages important for secondary cartilage induction and maintenance, and paralysis has a significant but differential effect on transcription of some of these genes. We find that *Fgf4* and *Fgf8* are dramatically affected by paralysis, indicating that their expression may be mediated by mechanical stimulation. Furthermore, FGF signaling activity is decreased following paralysis as indicated by the relative down regulation of *Pea3* and *Erm* transcription. While the role of FGF signaling in the context of cartilage, bone, muscle, and limb tendon is well described (Brent et al., 2005; Edom-Vovard et al., 2001; Eloy-Trinquet et al., 2009; Murakami et al., 2000; Ornitz and Marie, 2015), the influence of the mechanical environment on FGF signaling has remained unclear. While we do not observe an effect of paralysis on the transcription of TGFβ ligands or receptors, the downstream effector *Pai1* was significantly increased by paralysis, suggesting tissue atrophy and fibrosis in response to disuse (Naderi et al., 2009). There is a relationship between the mechanical environment and TGFβ signaling (Kleinnulend et al., 1995; Nguyen et al., 2013; Robbins et al., 1997; Shi et al., 2011), but how mechanical cues exert control over TGFβ signaling is not as well understood. Our results suggest that, in this context, TGFβ signaling activity is primarily regulated by post-transcriptional modifications like phosphorylation of SMADs (Anthwal et al., 2008; Berthet et al., 2013; Maeda et al., 2011; Wipff et al., 2007) and regulation of free, active TGFβ ligands, something we plan to pursue in future studies.

Knockouts of *Tgfβ2* and *Tgfβr2* in mice produce malformations of the dentary and its coronoid, condylar, and angular processes (Oka et al., 2008; Oka et al., 2007; Sanford et al., 1997), although, the malformations of the three processes likely arise via different mechanisms. Also, unlike duck and humans, the mouse coronoid process does not form via a secondary cartilage intermediate. In *Tgfβ2* null mice, the condylar and angular processes are smaller, but the secondary cartilages on these processes persist. However, secondary chondrogenesis was prevented by *Tgfβr2* knockout. Mandible culture experiments in mice also demonstrate that TGFβ signaling is required for condylar and angular secondary cartilage induction (Anthwal et al., 2008). In the context of our experiments, TGFβ inhibition does not produce bone defects, nor do we observe abnormalities in Meckel’s. This is consistent with TGFβ knockout data in which tendon formation is severely inhibited in the absence of *Tgfβ2, Tgfβ3*, or *Tgfβr2*, while primary cartilage is largely unperturbed (Pryce et al., 2009).

Our efforts to rescue paralyzed embryos led to the formation of a dense fibrous capsule and even cartilage around the bead. Although ligands were delivered using spherical beads and presumably diffused uniformly (Eichele et al., 1984), the axis of Alcian blue or Safranin-O positive tissue surrounding the beads is not radially symmetrical. Directional distribution of induced cartilage in quail and duck suggests that the mesenchyme and surrounding connective tissues overlying the surangular are not all equivalent in their capacity to generate secondary cartilage. Furthermore, the locations where cartilage is induced are spatially restricted to the general region where secondary cartilage forms in controls. Such a spatial constraint parallels published explant data in which the murine CP, which does not ordinarily form a secondary cartilage, can be induced to do so by fetal bovine serum (FBS)(Anthwal et al., 2015). Though FBS bathed the entire mandible, ectopic cartilage was only observed on the CP. In duck and quail, beads implanted too distal from the jaw joint, or too superficial, superior, or inferior to the surangular did not elicit a chondrogenic response.

Other experiments on developing limb tendons corroborate the ability of exogenous FGF and TGFβ ligands to maintain *Scx* even in the absence of mechanical stimulation, but to our knowledge, no instances of induced cartilage have been reported in those contexts (Edom-Vovard et al., 2002; Havis et al., 2016). The FGF and TGFβ signaling-dependent chondrogenic response we observed may be localized to tendon and connective tissues surrounding the MA insertion and is conserved between quail and duck. Though quail do not normally form secondary cartilage on their CP, the surrounding connective tissues are able to do so given the proper signaling environment.

Induced cartilage appears to be encapsulated and distinct from the surangular, mirroring native secondary cartilage development on the duck CP. Thus, the secondary cartilage on the CP is likely derived from cells in the tendon and adjacent connective tissue, not the periosteum as in articular secondary cartilage (Buxton et al., 2003). Experiments in other contexts suggest the existence of progenitor cells that express both tendon (e.g., *Scx, Tcf4*) and cartilage (e.g., *Sox9*) tissue markers that contribute functionally to establishing certain sites where tendons or ligaments insert onto primary cartilage and that such markers are involved in the patterning of these insertions (Blitz et al., 2013; Kardon, 1998; Kardon et al., 2003; Mathew et al., 2011; Schweitzer et al., 2001; Sugimoto et al., 2013). Cells that give rise to secondary cartilage on the CP may express a similar complement of lineage markers, which is supported by our previous expression analyses (Solem et al., 2011; Tokita and Schneider, 2009).

### Mechanical cues differentially regulate members of the FGF and TGFβ pathways

Clearly, musculoskeletal development and homeostasis depend upon proper biomechanical cues, however, the cell-biology that mediates this mechanosensation is not well understood. A variety of mechanisms including the primary cilium, Wnt signaling, and especially sclerostin, which is an osteocyte-specific Wnt inhibitor, have been implicated in mechanosensitive bone remodeling (Robling et al., 2016; Robling et al., 2008; Rolfe et al., 2014; Tu et al., 2012). Other potential mechanisms may include ligands being freed from the extracellular matrix, ion channels, focal adhesions, cytoskeletal dynamics, and many others (del Rio et al., 2009; Dupont et al., 2011; Hamill and McBride, 1996; Maeda et al., 2011; Mammoto and Ingber, 2010; Matthews et al., 2006; McBeath et al., 2004; Pruitt et al., 2014; Quinn et al., 2002; Raizman et al., 2010; Ramage et al., 2009; Roberts et al., 2001; Shakibaei and Mobasheri, 2003; Solem et al., 2011; Vincent et al., 2002; Vincent et al., 2007; Wang et al., 2009; Wen et al., 2017).

From our qualitative and quantitative analyses, a subset of genes stands out as likely mediating development of the MA complex (*Tgfβ2, Tgfβ3, Fgfr1*, and *Fgfr2*) as their abundance changes significantly and in the same direction regardless of whether the embryo was paralyzed or not (Fig.8A). This group of genes includes *Tgfβ2* and *Tgfβ3*, which induce chondrogenesis when delivered as ligands to paralyzed duck embryos or normal developing quail, suggesting that TGFβ signaling activity may be modulated post-transcriptionally and depend upon the availability of free, active TGFβ ligands. Also, we observed no change in *Tgfβr1, Tgfβr2, Tgfβr3, Smad3*, or *Smad7b.* Our analyses did find that one component of the TGFβ pathway is significantly more abundant in paralyzed samples. *Pai1*, a common transcriptional readout of TGFβ signaling (Kawarada et al., 2016), became significantly more abundant following paralysis. Our data support the hypothesis that TGFβ pathway-mediated responses to mechanical stimulation utilize post-transcriptional mechanisms. Quantifying free, active TGFβ ligands, or assaying phospho-SMAD abundance or nuclear localization would shed light on this phenomenon, something that we are working towards for future studies.

**Fig.8.**
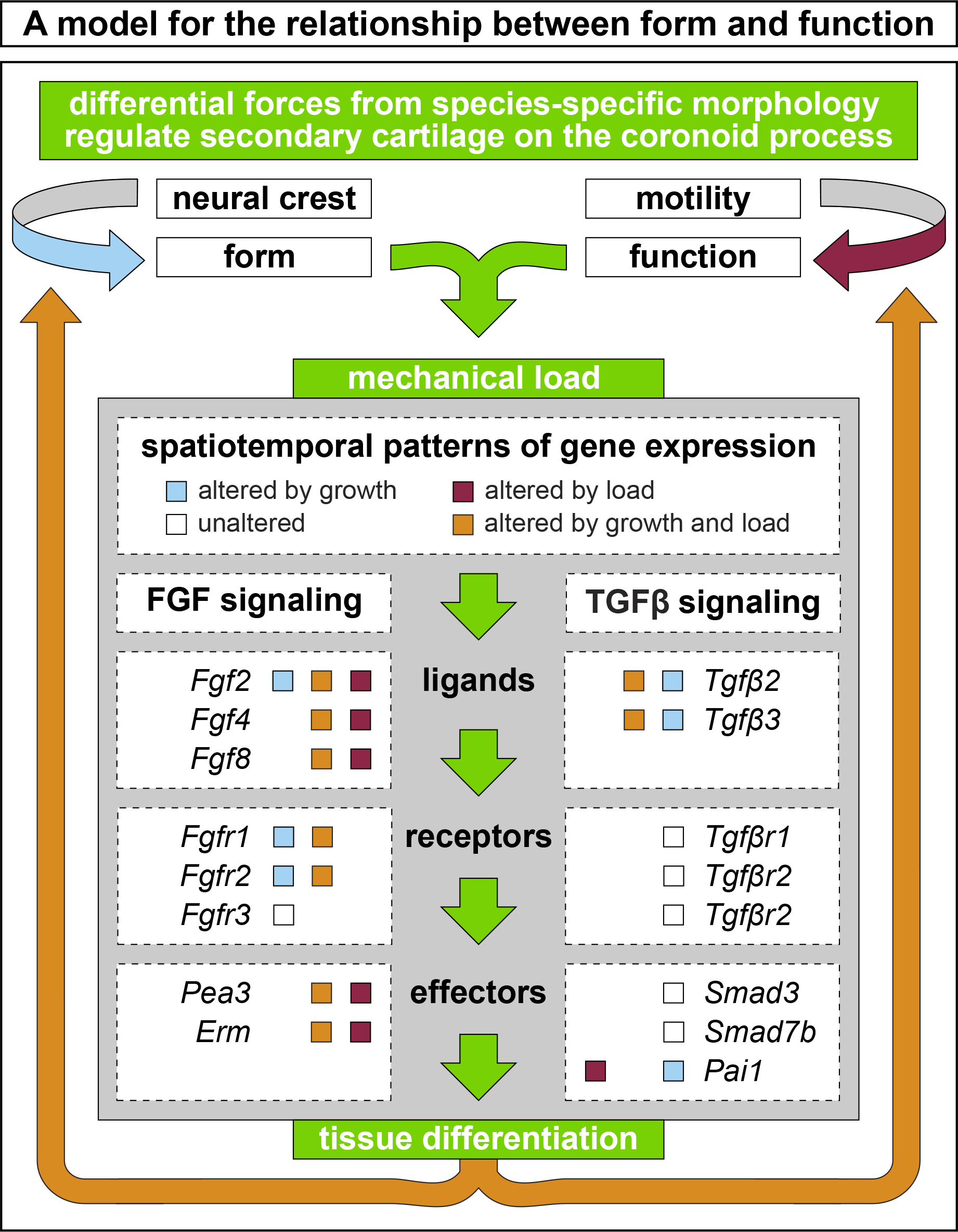
A model integrating form and function with FGF and TGFβ signaling. NCM-mediated species-specific jaw geometry, (i.e., dorsal versus lateral MA insertions) and functional loading by embryonic motility contribute to differential forces and tissue differentiation. The resultant mechanical stress leads to differential activation of FGF and TGFβ signaling and regulates the presence or absence of secondary cartilage on the CP. We observe three overlapping patterns of expression: One set is altered by growth (blue boxes), another altered by load (red boxes), and a third is altered by both growth and load (orange boxes). A fourth set of genes remains unaltered both during growth and despite a loss of embryonic motility (white boxes). Some genes are found in multiple sets, reflecting the complex integration of form and function during embryonic development.

Our analyses also indicate that a second set of five FGF signaling pathway components (*Fgf2, Fgf4, Fgf8, Pea3*, and *Erm*) likely mediates normal development of secondary cartilage and depends upon embryonic muscle contractions to maintain their activation. FGF signaling has been implicated in other mechanosensitive processes (Vincent et al., 2002; Vincent et al., 2007; Wen et al., 2017), but there is still a lot to learn about how FGF ligands, receptors, and transcriptional effectors interact with the mechanical environment.

Our data suggest a model (Fig.8) whereby species-specific secondary chondrogenesis on the CP arises as a consequence of functional motility acting upon NCM-derived form. In our model, the resulting stress within the insertion of the MA muscle onto the surangular differentially activates FGF and TGFβ signaling, which are each necessary and sufficient to induce chondrogenesis. Thus, by balancing cell-autonomous developmental programs and adapting to environmental cues, NCM generates species-specific jaw geometry and promotes structural and functional integration of the musculoskeletal system during development.

E.S. Russell in his classic book, *Form and Function* (1916) poses the question, “Is function the mechanical result of form, or is form merely the manifestation of function or activity? What is the essence of life, organisation or activity? (p.v)” Our findings provide evidence that form initially dictates function but then function modulates form. Cranial NCM establishes species-specific “organisation” prior to the onset of muscle “activity.” However, the musculoskeleton is developmentally plastic. As jaw activity begins, form adapts to meet and support functional demands. In the case of a duck, species-specific form, coupled with jaw activity, creates stresses within the MA insertion, differentially activates FGF and TGFβ signaling, and induces secondary cartilage on the CP. Appreciating the inextricable connection between form and function allows for a new perspective on the role of NCM in establishing form but also shows how the organism can modify that form to accommodate functional demands throughout development, under selective pressure, or in disease states.

## Acknowledgements

We thank J. Lotz, T. Alliston, R. Marcucio, J. Fish, and members of the Schneider lab for helpful discussions; Z. Vavrusova, M. Bodendorfer (Hague), D. Jaul, M. Chung, S. Smith, and D. Chu for technical assistance; and T. Dam at AA Lab Eggs for quail and duck eggs. This work was supported in part by NICHD T32 HD007470 and F31 DE024405 to K.C.W.; and NIDCR R01 DE016402, R01 DE025668, and S10 OD021664 to R.A.S.

## Author Contributions

R.A.S. and K.C.W. conceived of the project and designed the experiments; K.C.W. S.G., and S.H. performed the experiments; K.C.W. S.G., S.H., A.F., and R.A.S. analyzed the data; and R.A.S. and K.C.W. co-wrote the manuscript.

